# Embryonic circular RNAs at tandem duplicated genes in zebrafish present new paradigm in gene expression

**DOI:** 10.1101/220160

**Authors:** Vanessa Chong-Morrison, Tatjana Sauka-Spengler

**Affiliations:** University of Oxford, Weatherall Institute of Molecular Medicine, Radcliffe Department of Medicine, Oxford OX3 9DS, United Kingdom

## Abstract

Circular RNAs are a puzzling class of RNAs with covalently closed loop structure, lacking 5’ and 3’ ends. Present in all cells, no unifying theme has emerged regarding their function. Here, we use transcriptional data obtained by *biotagging* in zebrafish to uncover a high-resolution cohort of embryonic circRNAs expressed in nuclear and polysomal subcellular compartments in three developmental cell types. The ample presence of embryonic circRNAs on polysomes contradicts previous reports suggesting predominant nuclear localisation. We uncover a novel class of circRNAs, significantly enriched at tandem duplicated genes. Using newly-developed NGS-based approach, we simultaneously resolve the full sequence of putative "tandem-circRNAs" and their underlying mRNAs. These form by long-range *cis*-splicing events often between distant tandem duplicated genes, resulting in chimaeric mRNA transcripts and circRNAs containing their supernumerary excised exons, integrated from multiple tandem loci. Taken together, our results suggest that circularisation events rather than circRNAs themselves are functionally important.

---

Circular RNAs (circRNAs) are transcripts with an unusual structural organisation. Unlike linear mRNA transcripts, circRNAs have the positions of their 3’ and 5’ sequences swapped and are potentially formed via co‐ or post-transcriptional processes to produce characteristic backspliced circular transcripts^1^. Largely overlooked in the beginning, interest in circRNAs recently experienced a resurgence following genome-wide studies that demonstrated their ubiquitous nature and dominant presence in the human genome^2^ as well as in other organisms^3,4,5,6^. The biogenesis of circRNAs has been shown to be modulated by circularisation “signals” such as complementary sequences and/or secondary structures, which are embedded within the introns that flank circularised exons^7,8,9^. In terms of circRNA function, it has been suggested that they may function as miRNA sponges^10,3,11^, a finding further supported by a recent study of circRNAs in adult zebrafish^12^. CircRNAs can also regulate transcription of their parental genes in the nucleus^13^, and, in certain cases, may be translated into proteins^14^. Here, we identify an ensemble of putative circRNAs expressed at two different stages of zebrafish embryonic development, in three different cell populations from two subcellular compartments (nuclei and polysomes). We discover a novel cohort of nuclear circRNAs, enriched at tandem duplicated genes in the zebrafish genome (“tandem-circRNAs”). Although it is already known that the zebrafish genome has undergone a significantly higher rate of tandem duplications compared to other teleost species^15^, this phenomenon and its secondary regulatory effects on gene expression has until recently only been described in *Drosophila* ^16,17,18^ and in mammals^19^. Using our newly-developed screen-and-sequence circRNA pipeline, we describe and validate two novel “tandem-circRNAs”, *circ-cyt1* and *circ-smyhc,* and show that the splicing events across neighbouring tandem duplicated genes leading to their circularisation are correlative of the formation of chimaeric linear transcripts between the same tandem duplicated genes, echoing a previous observation in *Drosophila yakuba*^18^. Our results not only present additional implications in the elucidation of circRNA function, but also in the understanding of gene expression within the context of tandem duplicated genes and splicing mechanisms during transcription.

## Results and Discussion

### Identification of developmentally-regulated circRNA loci in zebrafish

We exploited previously obtained datasets comprising multiple ribo-depleted RNA-seq samples collected from two subcellular compartments - nuclear and polysomal, across three different cell types - migrating neural crest (*sox10*-expressing), myocardium (*myl7*-expressing), and cytoskeleton (*βactin2*-expressing; not shown) and at two developmental stages (16ss and 24-26hpf) (Fig. 1A)^20^. Putative circRNA candidates were identified at a rate of ∽6-12 circRNAs per million reads within each cell type (Table 1) using the published *find_circ.py* Python script^3^. Unlike in previous studies, which used whole-cell lysates, the analysis of total RNA transcripts obtained from subcellular compartments is poised to provide higher sensitivity for detecting putative circRNAs and yield new insights into circRNA biogenesis and cellular trafficking, as our current understanding of these processes is still in its infancy.

To identify tissue-specific clusters of circRNAs only present in either *sox10* or *myl7* datasets and not in their ubiquitous (*βactin2*) stage-matched control samples, we first performed the analysis without taking subcellular compartments into account (i.e. by pooling nuclear and polysomal data for *sox10* and *βactin2*). We recovered 125 and 373 putative circRNAs clusters which were uniquely produced in *myl7*- and *sox10*-expressing cells, respectively, while 112 and 153 clusters were exclusive to their stage-matched *βactin2* controls (Fig. 1B). The detection of circRNA loci was also developmental-stage-specific, as we identified 211 and 123 clusters produced in *βactin2*-expressing cells at 16ss and 24hpf stage, respectively (Fig.1B’). Finally, comparing the two divergent developmental cell types (*sox10*- and *myl7*-positive populations), in addition to 104 shared predicted circRNA clusters, we identified a relatively large number of circRNA candidates which were specific to either neural crest cells (*sox10,* 418) or to myocardium (*myl7,* 123) (Fig.1B”). Taken together, these results suggested that our putative circRNA loci were developmentally-regulated in time and space.

**Table 1.** Identification of putative circRNAs in *biotagging*^20^ RNA-seq datasets.

| Sample | Average no. of reads<br>(2 replicates/sample) | No. of circRNAs | % circRNAs predicted | circRNAs per<br>million reads |
| --- | --- | --- | --- | --- |
| *sox10* 16ss nuclear | 51,151,788 | 348 | 0.0007 | 6.803 |
| *βactin2* 16ss nuclear | 31,511,738 | 242 | 0.0008 | 7.680 |
| *myl7* 26hpf nuclear | 49,748,086 | 382 | 0.0008 | 7.679 |
| *βactin2* 24hpf nuclear | 34,220,784 | 337 | 0.0010 | 9.848 |
| *sox10* 16ss ribo | 46,661,325 | 556 | 0.0012 | 11.916 |
| *βactin2* 16ss ribo | 42,974,211 | 356 | 0.0008 | 8.284 |

We next sought to establish whether the developmental regulation of circRNA production was linked to the regulation and expression of their underlying linear mRNA transcripts. To test the relationship between the circRNAs and their corresponding mRNAs, we intersected the total read counts for each circRNA cluster in a given cell type with the total FPKM values of corresponding Ensembl-annotated gene(s) transcribed on the same strand. Where more than one mRNA transcript was associated with the circRNA cluster, we plotted the sum of the corresponding FPKM values in function of the total number of circRNA reads per locus (Fig.1C, C’). A weak positive correlation between the two types of transcripts (Pearson correlation of *ρ* = 0.297, 0.397, 0.397 and 0.447 for *sox10 16ss, βactin2 16ss, myl7 26hpf,* and *βactin2 24hpf,* respectively, Fig.1C) suggested that cell type-specific expression of circRNAs may be linked to the specific expression of their underlying genes, rather than being independently developmentally-regulated.

Finally, we used inverse PCR followed by Sanger sequencing to validate two circRNA candidates, *circ-fuca1* and *circ-gbgt1l4,* identified in *sox10* nuclear samples. Our results demonstrated the presence of ‘inverted’ back-splicing events thus suggesting that the predicted candidates were indeed circularised (Fig.1D).

### Putative zebrafish circular RNAs form genomic clusters and are compartment-specific

Next, we interrogated the identified circRNA loci to obtain individual circRNAs within each cluster (Fig.2A). Predicted circRNAs “congregated” within genomic loci overlapping annotated genes, with high prevalence for regions containing annotated Ensembl genes, rather than at gene-free domains. Of the total number of circRNA loci predicted per cell type; 76%, 79%, 80%, and 77% were found to overlap Ensembl annotated transcripts for *sox10 16ss, βactin2 16ss, myl7 26hpf,* and *βactin2 24hpf,* respectively. Moreover, we noted that circRNA clusters tend to contain more than one individual circRNA and have thus quantified the occurrences of putative circRNAs within the clusters. Identified putative circRNAs formed clusters consisting of 1 to 25

**Figure 1.**
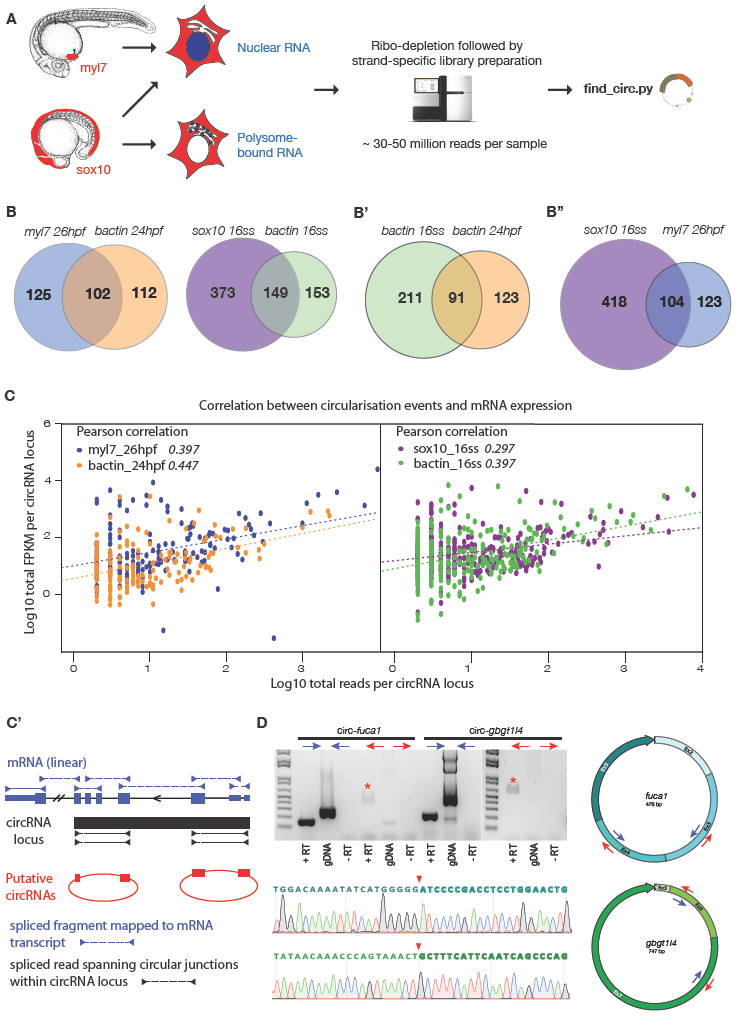
Putative zebrafish circRNA loci are developmentally-regulated and can be detected by inverse PCR (legend on next page).

A Workflow for identification of putative circRNAs during zebrafish embryonic development. **B** Venn diagrams showing overlapping and mutually-exclusive putative circRNA loci from each dataset according to cell type compared to ubiquitous control, **B’** at different developmental stages and **B”** from diverged cell types (*sox10* vs *myl7*). **C** Scatterplot comparison demonstrating weak correlation between expression of putative circRNAs and their underlying mRNA expression, where **C’** total read counts from each genomic cluster of circRNAs according to cell type were plotted against the total FPKM values of corresponding Ensembl-annotated mRNA transcripts found on the same strand. If more than one mRNA transcripts were annotated, the sum of FPKM values was used. **D** Validation of two *sox10* nuclear circRNAs by inverse PCR followed by Sanger sequencing. Blue arrows on gel image denote control non-inverse primer pairs and red arrows represent inverse primer pairs. PCR product indicative of circRNA shown with a red asterisk. Red solid arrows on Sanger sequencing traces show back-spliced junctions of *circ-fucaI* (top) and *circ-gbgt1l4* (bottom).

circularisation events within a particular genomic locus with the corresponding histograms showing non-normal, left-skewed frequency distribution of circRNA events per cluster. This demonstrates that, collectively, majority of circRNA clusters contained multiple rather than single putative circRNAs (Fig.2B). Regardless of the nature of circRNAs being quantified (loci/cluster vs individual candidates), our putative circRNAs appeared to be spatio-temporally regulated, with clear overlapping as well as mutually exclusive groups (Fig.2C). Full lists of individual circRNAs for each dataset are available as Supplementary Material.

We next surveyed our subcellular compartment-specific datasets to identify circRNA content separately in isolated nuclei and polysomes for *sox10*- and *βactin2*-expressing cells at 16ss. Surprisingly, we found that in addition to the nuclear compartment, individual circRNAs localised within the polysomal compartment in both cellular contexts studied. The identified circRNAs were either common to both or exclusive to either compartment (Fig.2D) and could also be detected via poly(A)-selection of the polysomal RNA pool in *sox10*-expressing cell populations (Supp.Fig.1A), suggesting that circRNAs may remain associated with their cognate mature mRNAs. Our finding that circRNAs were associated with polysomes (Fig.2D) contradicted previous candidate approach-based studies that suggested circRNA(s)-of-interest were always depleted in the polysomal-bound fraction^1,6^, and functioned either in the nucleus^13^ or as miRNA sponges in the cytoplasm^3,10^. However, our findings were in line with recent ribosome profiling studies that demonstrated the association of non-coding RNAs (including 5’UTRs) with ribosomes, suggesting that presence at the polysomal compartment does not necessarily equate to translation into proteins^21^. Furthermore, recent evidence has suggested that certain circRNAs may be translated into proteins^14^. Detected polysomal circRNAs were also discovered in a tissue-specific fashion, i.e. identified either in *sox10*- or *βactin2*-expressing cell populations or shared between the two cell types, suggesting that they also could be developmentally-regulated (Supp.Fig.1B).

**Figure 2.**
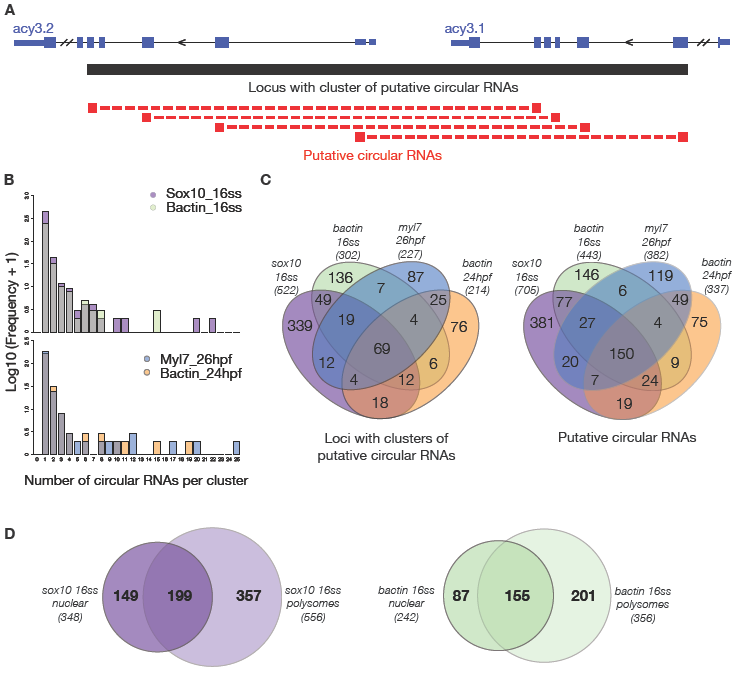
Putative zebrafish circular RNAs form genomic clusters and are compartment-specific.

A Schematic representation of a genomic locus (*acy3.1/2*) with multiple overlapping circularisation events predicted. B Histogram shows frequency distribution of number of circRNAs predicted per genomic cluster of circRNAs (from 1 to 25 circRNAs/cluster), according to cell-type. C Venn diagrams show overlapping as well as mutually-exclusive genomic clusters of putative circRNAs (left) and individual putative circRNAs (right) according to cell type (*sox10 16ss* and *βactin2 16ss* consist of both nuclear and polysomal compartments, *myl7 26hpf* and *βactin2 24hpf* consist only of nuclear compartment). D Venn diagram showing overlapping and mutually-exclusive putative circRNAs from each subcellular compartment (nuclear or polysomal) from *sox10 16ss* and *βactin2 16ss.*

### CircRNA “hotspots” in the nuclear compartment are enriched at tandem duplicated genes

Putative circRNAs identified in our study had up to ∽50-fold lower abundance compared to previous studies^3^’^6,12^, likely reflecting differences in species, RNA preparation, cell types, choice of algorithm software for circRNA prediction^22^, or the nature of our samples. We speculated the last scenario to be the most likely, as we used much smaller quantities of material isolated in a compartment‐ and tissue-specific fashion from the *in vivo* context of developing embryos instead of abundant whole cell lysates from large quantities of cultured cells or whole tissue from embryos. With this in mind, we selected circRNAs detected across multiple datasets to generate a high-confidence set of circRNAs for validation, reasoning that these circRNAs were more likely to be functional and be of potential biological significance. We thus chose to scrutinise the circularisation events that were present in four nuclear datasets encompassing an extensive complement of cell types and developmental stages - *sox10* (16ss), *βactin2* (16ss), *βactin2* (24hpf) and *myl7* (26hpf), and defined this set as circRNA “hotspots” (Fig.3A; see Supplementary Material (Tables)).

Our finding that multiple-circRNA clusters occurred with high frequency (Fig.2B) suggested a general feature of our circRNAs where they may be preferentially formed in particular regions of the zebrafish genome. Upon close examination, we noted that many of our “hotspot” circRNAs overlapped tandem duplicated genes in the zebrafish genome^23^. Recent genome duplication events that took place early in the teleost lineage before its radiation resulted in a remarkably large number of duplicated genes, particularly in the zebrafish genome which has a more significant proportion of tandem versus intra-chromosomal duplicated genes compared to other teleost fish^15^. Moreover, the predicted circularisation junctions of individual circRNAs mapping to a duplicated region were often separated and found within neighbouring duplicated loci rather than in a single gene, thus suggesting that circRNA junctions could span several kilobases between tandem duplicated genes. We quantified this observation and found that “hotspot” circRNAs overlapping duplicated genes, a subset which we now class as “tandem-circRNAs”, were highly significantly enriched in our high-confidence “hotspot” set of circRNAs (56 out of 112 “hotspots”; hypergeometric *p*-value of 6.31 x 10^-11^) (see Supplementary Material (Tables)). To explore co‐ or post-transcriptional splicing events that underlie tandem-circRNAs, we performed *de novo* transcript assembly for each dataset using StringTie^24^. Interestingly, for 20 out of 56 putative tandem-circRNAs, we detected one or more “chimaeric” fusion transcripts that included exons from multiple (two or more) duplicated neighbouring loci with unannotated splicing junctions to link one tandem transcript to another, in addition to successful assembly of transcripts consisting of exons from individual tandem duplicated genes with predicted splicing junctions (Fig.3A’).

We next sought to validate the presence of tandem-circRNAs and resolve the full composition of the circRNA transcripts. Current biochemical methods are mainly aimed at proving the circularity of putative circRNAs and include RT-PCR followed by inverse PCR, Northern Blotting with probes designed to target the circularised junction sequences, and RNaseR sensitivity tests^25^. Another validation method takes advantage of the high processivity and strand displacement capabilities of reverse transcriptases which can yield cDNAs consisting of concatemers of circRNAs, a phenomenon known as rolling-circle RT. When this phenomenon is observed, characterised by a laddering appearance of PCR products on a gel, it provides substantial evidence for circularisation^6^.

**Figure 3.**
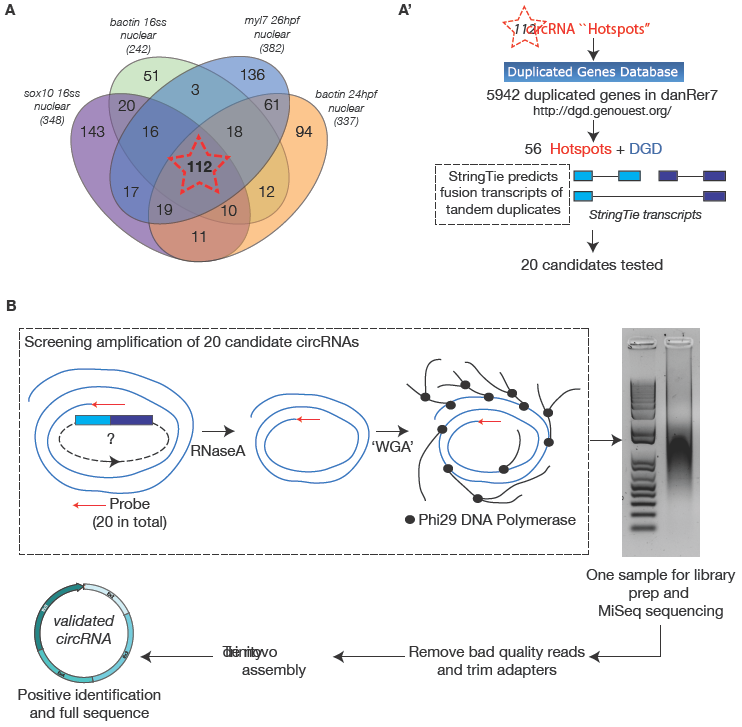
Validating circRNA “hotspots” in the nuclear compartment enriched at tandem duplicated genes.

**A** Venn diagram shows overlapping and mutually-exclusive putative circRNAs from nuclear compartments of each cell type with their corresponding developmental stages. **A’** 112 putative circRNAs found in all datasets (“hotspots”, red star) were examined and cross-checked with 5942 zebrafish duplicated genes from the Duplicated Genes Database. 56 “hotspots” were found to overlap regions of duplicated genes. 20 high-confidence candidates were picked based on underlying splicing events between tandem duplicated genes (predicted and assembled using StringTie *de novo* assembly) that may indicate circularisation events between the linear transcripts. **B** A novel “screening amplification” pipeline (see Methods) was established to simultaneously probe for circRNA expression and resolve the full sequence of circRNAs detected. CircRNA(s) were reverse-transcribed, their cDNA purified and amplified using a Whole Genome Amplification-based (‘WGA’) method.

Here, we developed a new screen-and-sequence method that allows multiplexed testing of up to 20 putative circRNAs in a single reaction. Briefly, half of an RNA sample was first enriched by ribo-depletion to remove rRNA species but still include circRNAs, and the remaining half was poly(A)-selected to mostly exclude circRNAs and to focus on mRNA transcripts. Both samples were then reverse transcribed with a highly processive reverse transcriptase using a primer set of 20 oligo DNA probes (see Supplementary Material (Tables)) designed as reverse-complement primers to the exact circularised junctions-of-interest from 9 different duplicated gene loci containing “hotspot” circRNAs. The reverse transcription was allowed to proceed in conditions conducive to rolling-circle RT (see Methods). RNaseA treatment was performed to remove residual RNA after reverse transcription was complete. cDNA was purified and concentrated for a primer-free unbiased whole-genome amplification approach^26^. The amplified products were sequenced using Illumina MiSeq Platform and a bioinformatics pipeline utilising *de novo* transcript assembly and BLAT was used to extract original circRNA template sequences (Fig.3B) (see Methods). In parallel, we performed inverse PCR of the same 20 candidates, where Tapestation profiles of the purified successful PCR reactions also demonstrated periodicity in the product lengths indicative of rolling-circle RT (Supp.Fig.2).

We present an in depth analysis of two assembled transcripts, c119_g1_i1 (c119) and c408_g1_i1 (c408), identified using our screen-and-sequence validation approach and found to overlap annotated duplicated gene regions at chromosome locations chr19:5,975,453-5,987,999 (DGD_ID Group_1158) and chr24:42,242,905-42,325,671 (DGD_ID Group_1489), respectively^23^. Both transcripts were only detected in the ribo-depleted amplified sample and not in the poly(A)-selected amplified sample (Fig.4A and Supp.Fig.3A; 1st and 2nd track). The putative full length circRNA transcript c119 at DGD_ID Group_1158 (Fig.4A; 5th track from the top in maroon) was identified using the single oligo DNA probe (that was part of the 20-oligo probe pool) targeting a circular junction between the 3rd exon (splice acceptor) of the *cyt1* gene and the 2nd exon (splice donor) of the *zgc:109868* gene positioned upstream of it, within the region harbouring the predicted “hotspot” tandem-circRNA junction being tested (Fig.4A; blue track). Transcript c119 contained a back-spliced sequence, with full or partial exons from one tandem gene to another in this locus (Fig.4A; pink dotted box). Despite high sequence similarity between tandem duplicated genes (demonstrated by multiple mapping of transcript c119), we were able to obtain the full sequence of the circRNA (522 bp), as the extracted sequence of transcript c119 clearly presented a periodicity indicative of rolling-circle RT. This periodically repeating pattern of the now-identified circRNA, unique sequences within exon II (from *zgc:109868*) and exon III (from *cyt1*), and a unique alternative back-splicing junction between partial exon IV and exon I allowed us to eliminate the ambiguity due to sequence identity in exon I and exon IV across the tandem genes within the locus (*cyt1, cyt1l* and zgc:109868) thus assigning the partial exon IV and full exon I to *cyt1* and/or *cyt1l* genes, respectively. Altogether, these observations presented strong evidence for the identified tandem-circRNA, which we designated as *circ-cyt1* (Fig.4B). *Kallisto*^27^ quantification estimated the abundance of *circ-cyt1* at ∽158 Transcript Per Million (TPM), calculated based on the periodicity, from transcript c119’s abundance at 330 TPM. The alignment of the circRNA junction and *de novo* assembled StringTie transcripts spanning the tandem duplicated genes (Fig.4A; 4th track from the top in red) demonstrated that most of the exons annotated within the StringTie transcripts were not present in the assembly. The positions of these “lost/excised” exons were matched and demarcated by the circRNA backsplicing event.

Based on this “complementary splicing” observation, we further hypothesised that the *de novo* assembled transcripts represented linear RNA transcripts formed as a result of circularisation events between the tandem duplicated genes, with a role to remove supernumerary exons as a result of duplication events and still generate functional protein. As such, events of circularisation would offer a unique mechanism by which null-products of prolonged, erroneous transcription would be removed to generate a single copy of the duplicated tandem gene harbouring chimaeric exons from different duplicates. It is of note that highly conserved sequence content is most likely essential for enabling the circularisation events, as it ensures the preservation of affinity between pairs of splice donors/acceptors to still preferentially be selected during back-splicing.

Transcript c408 at DGD_ID Group_1489 (Supp.Fig.3A; 5th track from the top in maroon) was identified using seven oligo DNA probes (that formed part of the 20-oligo probe pool) targeting different highly-supported circular junctions within this region. This domain contained a large cluster of predicted “hotspot” tandem-circRNAs, with circRNA junctions spanning the exons between multiple tandem duplicated genes (*smyhc1, smyhc2, smyhc3* and *si:ch211-24n20.3*) (Supp.Fig.3A; blue and red track). Similar to transcript c119, transcript c408 contained a back-spliced sequence, with full or partial exons from the *smyhc/si:ch211-24n20.3* locus (Supp.Fig.3A; pink dotted box). In this case, however, high sequence similarity between the tandem duplicated genes hindered definitive conclusion for half of the individual exonic “parts” that form the identified template. Nevertheless, transcript c408 also demonstrated a periodicity indicative of rolling-circle RT, thus presenting strong evidence for circularity of the identified circRNA. Its full sequence (292 bp), designated as *circ-smyhc,* was a chimaeric circular transcript containing back-spliced exons from *smyhc1* and most likely from up to four different genes (*smyhc1, smyhc2, smyhc3* and/or *si:ch211-24n20.3*) (Supp.Fig.3B). *Kallisto* quantification estimated the abundance of *circ-smyhc* at ∽79 TPM, calculated based on the periodicity, from transcript c408’s abundance at 162 TPM. Similar to the previous locus described, “complementary splicing” events within the underlying linear mRNA transcripts were also predicted. Full sequences of both *circ-cyt1* and *circ-smyhc* are provided as Supplementary Material (File).

**Figure 4.**
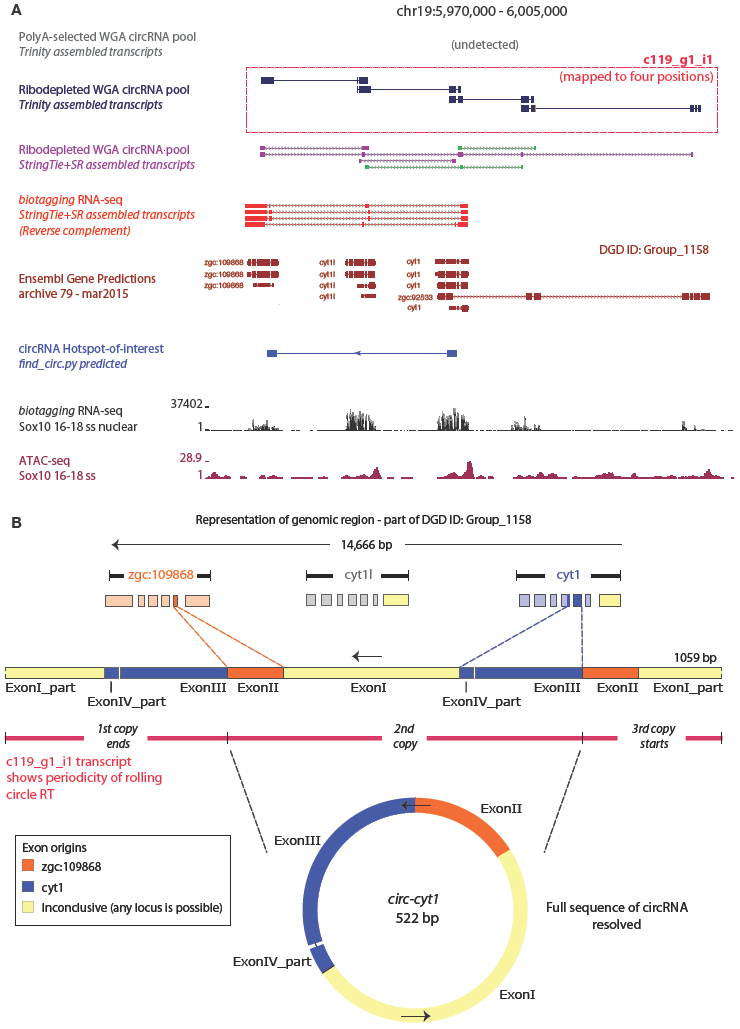
Identification and characterisation of *circ-cyt1* (legend on next page).

**A** Genome Browser snapshot of genomic region where the tested circRNA “hotspot”-of-interest resides. Trinity assembled transcript c119_g1_i1 (pink box) was detected only in the ribo-depleted amplified sample but not in the poly(A)-selected sample, and therefore its sequence was extracted for further characterisation. RNA-seq and ATAC-seq tracks provided information on overall gene expression levels and position of the promoters of each tandem duplicated gene. **B** Trinity transcript c119_g1_i1 demonstrated sequence periodicity indicative of rolling-circle reverse transcription, strongly suggesting circular nature of original RNA template. Full map of identified *circ-cyt1* is shown. Exons with inconclusive gene origins (due to sequence similarity between tandem duplicated genes) are coloured pale yellow.

To conclude, our results present a novel insight into the genomic organisation during circRNA biogenesis, within the context of zebrafish embryonic development. The enrichment of high-confidence nuclear compartment circRNAs at tandem duplicated genes suggests a previously unknown mechanism and role for tandem-circRNAs. Tandem duplicated genes are often tightly packed within a given genomic region and present a potential for producing long transcripts across multiple duplicated genes. We show that such closely-related tandem duplicated transcripts (but highly similar at the level of protein structure) undergo circularisation to generate back-spliced chimaeric circular transcripts, with contributing exons originating from multiple tandem genes, possibly to enable the production of a single functional full-length mRNA transcript per duplicated region and ensure translation of a correct protein. This idea is reminiscent of one of the earliest circRNA to be described, *circ-Sry. Circ-Sry* was found to be the most abundant *Sry* RNA molecule in the testis except within the genital ridge, where the linear *Sry* transcript was predominantly expressed and important for sex determination in mammals^1^. Capel *et al.* suggested that circular transcripts could represent an additional mechanism to prevent translation, similar to splicing variants that introduced alternative stop codons, as initially observed in *Drosophila,*^28^. Taken together, these findings revisit one of many enigmatic questions relating to our understanding of how and why “genes” are expressed, by highlighting innocuous but prevalent transcription of the genome for reasons other than to make any obvious protein.

## Methods

### circRNA identification

Initial circRNAs were identified using compartment-specific and cell type-specific transcriptional datasets obtained in our previous *biotagging* study^20^. Here we used the corresponding stranded sequenced RNA-seq libraries that were previously mapped and analysed for linear RNA content, to search for presence of embryonic circRNAs using the well-established pipeline, *find_circ.py*^3^. CircRNA datasets from biological replicates and/or same cell types (when pooling nuclear and polysomal compartments) were combined using bedtools intersect.

### Screen-and-sequence rolling-circle RT

Nuclei were collected using the *biotagging*^20^ method by crossing ubiquitious BirA driver to ubiquitious Avi effector. RNA was extracted using RNAqueous^TM^ Micro (AM1931, ThermoFisher Scientific). One half of RNA was ribo-depleted using RiboErase (KK8483, KAPA Biosystems) and the other half was poly(A)-selected using NEBNext^TM^ Poly(A) mRNA Magnetic Isolation Module (E7490, New England Biolabs). Reverse complement reads corresponding to predicted circular RNA junctions were extracted and aligned using Clustal Omega Multiple Sequence Alignment tool to obtain consensus DNA sequences for each junction-of-interest. Sequences were ordered as standard ssDNA oligos from Integrated DNA Technologies (IDT), and pooled to obtain a final concentration of 2pmolμL^-1^ per probe. Reverse transcription was performed in parallel for both ribo-depleted and poly(A) RNA samples using SunScript^TM^ Reverse Transcriptase RNaseH‐ (421010, Expedeon) with the following conditions; denaturation at 68 °C, followed by reverse transcription at 70 °C for 1 hour with additional spiking of dNTPs in the middle of the incubation period. Following rolling-circle RT, 0.2 μL of 10mg mL^-1^ RNaseA (EN0531, ThermoFisher Scientific) was added to the reactions and incubated at 37 ^°^C for 1 hour. The resulting cDNA pools were purified using Monarch^TM^ PCR & DNA Cleanup kit (T1030, New England Biolabs). Amplification was performed using TruePrime Whole Genome Amplification kit (370025, Expedeon) according to manufacturer’s protocol. Amplified products were purified by phenol/chloroform extraction followed by ethanol precipitation for library prepartion using Nextera DNA Library Prep kit (FC-121-1030, Illumina). Libraries were sequenced on the MiSeq platform using MiSeq v2 300 cycles kit (MS-102-2002, Illumina).

### Bioinformatics processing

Illumina adapters were trimmed from raw reads using *scythe* ^29^. Adapter-trimmed reads were further trimmed based on quality using *sickle*^30^ and the following parameters; `-t sanger`, `-l 50`, `-q 30`, to retain Q30 reads of at least 50bp in length. Using these processed reads as input, *de novo* transcript assembly was performed using Trinity^31^ with the following parameters; `-JM 100G` and `-no_bowtie`. Prior to BLAT alignment^32^, an over-occurring 11-mer tile of the *D. rerio* genome (danRer7) was created using `-makeOoc`. The Trinity output FASTA file containing all the assembled transcripts were aligned using BLAT default parameters. Alignment results were converted to BAM format and viewed on UCSC Genome Browser to identify Trinity-assembled transcripts that overlapped loci containing junctions-of-interest, in the ribo-depleted sample only. Full sequences of promising transcripts were imported into SnapGene software for further sequence analysis (e.g. sequence periodicity, exon/intron structure). To obtain possible mRNA splicing events between tandem duplicated genes at loci with putative circRNAs, an alternative *de novo* transcript assembly approach was used on *biotagging*^20^ RNA-seq datasets. Raw reads were processed as described above for Trinity, but assembled using the StringTie+SR approach^24^ with the following *superreads.pl* paramaters; `-s 1` and `-j 14100000000`. “LongReads” were mapped to the genome using HISAT^33^.

## Data Availability

The raw sequencing data used in this study can be accessed via Accession ID GSE89670, whereas all the processed data is available as a part of Supplemental Material and can also be downloaded from the ‘circRNAs’ repository on our GitHub account (https://github.com/tsslab/circRNAs).

## Acknowledgements

This work was supported by MRC (G0902418), Lister Institute Research Prize, Oxford BHF CRE award (RE/08/004) to T.S.-S. and a Clarendon Fund Fellowship to V.C.-M.

## Declaration

The authors declare no competing interests.

## Author contributions

Conceptualisation, methodology, analysis, and writing by V.C.-M. and T.S.-S. Funding acquisition by T.S-S.

